# Statistical Physics of DNA melting: Unveiling the artificial corrections for self-complementary sequences

**DOI:** 10.1101/2020.11.02.365098

**Authors:** C. A. Plata, S. Marni, A. Maritan, T. Bellini, S. Suweis

## Abstract

DNA hybridization is at the heart of countless biological and biotechnological processes. Its theoretical modeling played a crucial role, since it has enabled extracting the relevant thermodynamic parameters from systematic measurements of DNA melting curves. However, in its current state, hybridization modelling requires introducing an extra entropic contribution in self-complementary sequences that lacks any biophysical meaning. In this article, we propose a framework based on statistical physics to describe DNA hybridization and melting in an arbitrary mixture of DNA strands. In particular, we are able to analytically derive closed expressions of the system partition functions for any number *N* of strings, and explicitly calculate them in two paradigmatic situations: (i) a system made of self-complementary sequences and (ii) a system comprising two mutually complementary sequences. We derive the melting curve in the thermodynamic limit (*N* → ∞) of our description, which differs from the expression commonly used to evaluate the melting of self-complementary systems in that it does not require correcting terms. We provide a thorough study comprising limit cases and alternative approaches showing how our framework can give a comprehensive view of hybridization and melting phenomena.

**SIGNIFICANCE:** In this study, we provide a transparent derivation of the melting curves of DNA duplexes using basic tools of statistical mechanics. We find that in the case of self-complementary sequences, our expression differs from the one used in literature, which is generally amended by the introduction of a phenomenological correction which in our approach becomes unnecessary. By offering a clean formal description of DNA hybridization, our approach sharpens our understanding of DNA interactions and opens the way to study the pairing of DNA oligomers away from any thermodynamic limit.

## INTRODUCTION

The selective interaction and pair formation of nucleic acid polymers and oligomers is the basic mechanism enabling gene coding and replication. It also at the core of a wealth of other biological processes, such as gene regulation and secondary structuring of RNA, of biotechnologies, such as PCR and SELEX, and of DNA-based nanotechnologies and DNA origami. These interactions are based, for the largest part, on the Watson-Crick pairing of complementary bases (1). DNA and RNA pairing has a limited interval of stability. When submitted to enough stress, either physical (temperature, competing forces) or chemical (solvent composition) the double helix denaturates. In particular, thermal denaturation, leading to the unfolding of a single strand or to the splitting of a duplex, is of paramount importance in technologies, including amplification, screening and sequencing (2–6). Therein, tailored nucleic acids sequences usually work as probes unveiling biological information of a sample or mediating for amplification during polymerase chain reaction.

Thermal denaturation of duplexes provides an easy but crucial access into DNA and RNA thermodynamics. Melting curves, typically represented as plots of the fraction of paired DNA oligomers vs. temperature (T), can be experimentally accessed either via UV absorbance, based on the so-called hypochromicity effect (7–11), or by measuring the fluorescent emission of environment-sensitive DNA or RNA-binding fluorophores. In both cases, measurements directly yield, after suitable normalization, the melting curve. Melting curves are mainly characterized by their characteristic temperature, the so-called melting temperature *T_m_*, and by the sharpness of the bound-unbound transition, both depending on the energetics of pair interactions. While some very simple physical models, based for instance on the Ising model (12) or Haminoltonian mechanics (13), have been used to describe such interactions, the most accepted and used model describing the interaction of nucleic acid duplexes is the nearest-neighbor (NN) model. Originally introduced by Tinoco et al. in order to study the thermostability of RNA (14, 15), it has developed into a detailed protocol enabling to predict *T_m_* of strands of arbitrary length ans sequence. The model has been recently applied even under circumstances of molecular crowding (16).

According to the NN model, the free energy involved in the pairing of two strands depends on the specific nucleotide sequence and can be obtained as a sum of contributions stemming not just from individual base pairs, but from couples of nearest base pairs. Such a splitting incorporates the notion that the duplex stability is largely depending on stacking forces, acting between neighbouring molecular planes of paired bases. The computation of melting temperature *T_m_* in the NN model requires knowing the specific contributions to the free energy conveyed by each couple of nearest neighbour paired bases, which depends on the identity and location of the four nucleobases involved in such a segment of double helical structure. Conversely, the knowledge of the *T_m_* of a wide number duplexes of variable length and sequence enables determine of the thermodynamic parameters associated with each type of couples of nearest base pairs (17–26).

The extraction of such thermodynamic parameters necessarily involves a comparison between experimental and theoretical melting curves. The latter is typically based on a two-state transition model, which neglects any state intermediate between the intact duplex and the fully melted strands (27–29). The theoretical derivation of the melting curve stands on the calculation of either the system partition function or of the equilibrium constant in the balance equation of the two-state model. The functional shape of the melting curve slightly varies when considering complementary or self-complementary sequences. However, in the latter case, in order to have an agreement between theory and experiment, the introduction of an extra symmetric correction (19, 21, 22, 25, 26, 28) is required.

In spite of the success when reproducing experimental data, herein we submit to revision the current formulation of the theory of DNA melting curves. Specifically, our aim is twofold. On the one hand, we want to exploit the parallelism between the problem of DNA melting and the process of disassociation and recombination of diatomic molecules. From the physical perspective, they are equivalent problems and the latter has been very well studied in the past (30, 31). Specifically, solutions for the problem involving diatomic molecules have been obtained with exact mathematical treatment even for systems with a low number of components, away from the thermodynamic limit. On the other hand, we want to shed some light on the origin of the symmetric correction for self-complementary sequences, which is normally used in NN model computations, but lacks of a clean justification. As we will show, in our approach there is no need for such extra entropic contribution.

To achieve our goals, we put forward a clear formulation of the melting curve based on the partition function calculation. We present exact results for arbitrary DNA mixtures that, afterwards, are thoroughly analyzed in the thermodynamic limit for systems with experimental relevance.

## METHODS

### Partition function for a self-complementary mixture

We consider a system made by *N* identical self-complementary oligomers enclosed in a total volume *V* at temperature *T*. The system is characterized thus by a concentration *c* = *N/V*. Oligomers can be either free or paired, resulting in a duplex. Free oligomers give a contribution *G_f_* to the global Gibbs free energy, whereas each duplex contributes with *G_p_*. Henceforth, the free energy difference due to the inner degrees of freedom is Δ*G*_0_ = Δ*H*_0_ − *T*Δ*S*_0_ = *G_p_* − 2 *G_f_*. There is an extra entropic cost of the pairing, since paired oligomers cannot explore freely the full phase space. Specifically, we consider that pairing interaction has a finite range characterized by a volume *V*_0_, which formally means that if oligomers *i* and *j* are paired, then the relative position vector 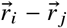 of their centers of mass is enclosed in a volume *V*_0_.

Since the system is in contact with a thermal bath at temperature *T*, the canonical distribution provides us with the probability *p n* of having *n* duplexes formed in the system,

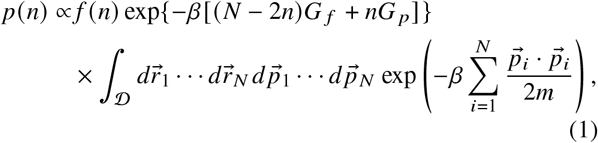

where 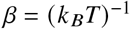, *k_B_* is the Boltzmann constant (which has to be substituted by the gas constant *R* when the free energy is expressed in molar quantities), 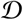 is the domain of the 6*N*-dimensional available phase space, and *f*(*n*) is the number of possible different configurations with exactly *n* duplexes formed in our system (with a total number of oligomers equal to *N*). Note that the thermodynamic potential of interest is the Gibbs free energy, which has been previously defined, since we consider that the number of oligomers, the pressure and the temperature is kept constant. Moreover, in this expression, we take into account the degeneracy of the state through *f*(*n*), whereas the integral and the first exponential account for the phase space explored by the centers of mass of the oligomers and the inner degrees of freedom, respectively. In the integral of Eq. (1), there are *N −* 2*n* free oligomers and 2*n* paired oligomers. Therefore,

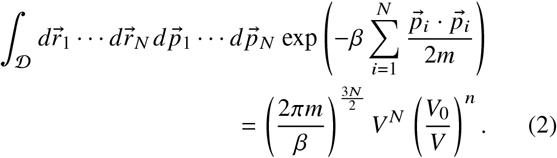

We thus get that the probability of having *n* couples of paired oligomers is

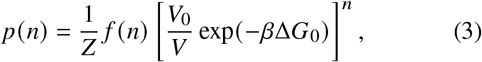

where we have absorbed conveniently some constant factors with respect to *n* in the partition function *Z*. It comes in handy to get rid the appearance of *V*_0_ from our expressions. To do that, we write it as an additive entropic contribution, defining the weighted Boltzmann factor

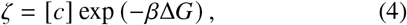

where [*c*] is the number concentration measured in mol/L and Δ*G* = Δ*H* − *T*Δ*S* is the global free energy, which includes the effect of *V*_0_, with Δ*H* = Δ*H*_0_ and Δ*S* = Δ*S*_0_ + *k_B_* with ln (*N_A_*[*V*_0_]) with *N_A_* being Avogadro’s number (the square brackets in *V*_0_ just indicates that it is consistently measured in L). Note that, usually, experimental measurements of the entropy of a duplex hardly distinguishes between the different contributions to the total entropy change, which makes our choice especially convenient. Importantly, the choice of units for the concentration and the volume has to be consistent. We have chosen the liter since this is the unit usually used in literature and consequently it leads to the values of Δ*S* reported in literature. The freedom to redefine a different factor before the exponential in Eq. 4 introducing an extra correcting contributions in Δ*S* will play an important role in our final results.

Using the Boltzmann factor introduced above, we obtain finally that the probability of observing *n* duplexes, or equivalently, the probability of being in the hybridization state *n*, is

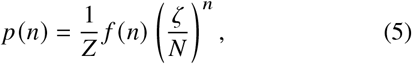

where the partition function guarantees the correct normalization, that is,

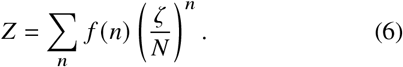

The sum above is defined over all possible values of *n*, which are the integers such that 0 ≤ 2*n* ≤ *N*.

At last, we need to write the degeneracy *f*(*n*) of an hybridization state

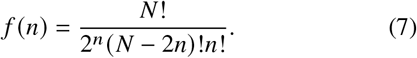

The factors in the denominator takes into account the different symmetries of a hybridization state under exchanging oligomers. Respectively, one can exchange oligomers within each duplex, all the free oligomers, and all the duplexes.

The partition function contains all the relevant statistical information which is useful to derive the expected values for the hybridization state. Specifically one can find straightforward that the average number of duplexes equals

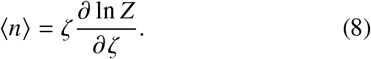

Later on, we will perform explicit computation of both the partition function and the average number of duplexes. Nevertheless, we focus first on studying a general formalism for analyzing arbitrary mixtures of oligomers.

### Partition function for an arbitrary mixture

In this subsection, we consider the most general system made by hybridizing oligomers. Specifically, we consider *S* different sequences labeled from 1 to *S*. We call the number of oligomers with the sequence *i* present in our system *N_i_*. Therefore, 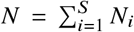 is the total number of oligomers in the mixture. Hence, the number concentration of the *i*-th sequence is *c_i_* = *N_i_*/*V*, whereas again the total concentration of oligomers is *c* = *N/V*.

We consider that, in principle, any duplex formation (complementary or not) may occur. In particular, we call the global free energy change Δ*G_i j_*, including the entropic contribution stemming from the corresponding *V*_0_, in a duplex formation comprising oligomers with sequences *i* and *j*. Consistently, we can define the Boltzmann factors

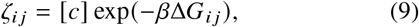

analogous to Eq. 4. Of course, in a real mixture, some attachments may be very unlikely, which means that such duplex has a very high value of Δ*G*, or equivalently a very low value of *ζ*.

In this general case, the hybridization state is not characterized just by one number. On the contrary, we need a vector 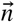 containing the number of all formed duplexes *n_i j_* comprising oligomers with sequences *i* and *j* in our system. Note that *n_i j_* = *n_ji_* by definition. In spite of this dimensional difference, the probability of having a certain hybridization state 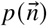 is provided by the canonical distribution

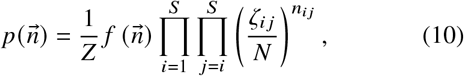

with the partition function

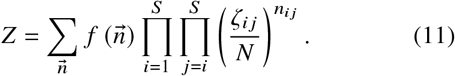

On this occasion, the sum over 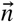 is defined over all possible values of *n_i j_* that are integers and holding the set of inequalities 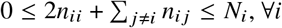. Fllowing the same reasoning that leaded to Eq. 7, the degeneracy 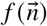 can be explicitly written as

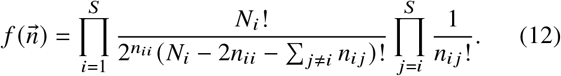

Again, the partition function contains all the relevant statistical information from which we can profit to derive expected values. In particular, the average number of duplexes comprising oligomers with sequences *i* and *j* is given by

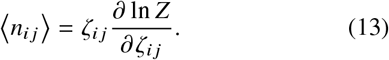

Note that all the theory converges consistently to the one developed in the previous subsection for self-complementary mixtures, when considering *S* = 1 and with the identification *n*_11_ = *n*.

We expect that a high level of hetereogenity in the mixture will lead to greater difficulties in the computation of the partition function and the successive physical quantities of interest. Remarkably, beside the self-complementary situation with *S* = 1, there is another case of particular interest that can be calculated: when two sequences that are complementary. In this instance, in fact, the chances of having an energetically convenient hybridization between two identical oligomers is very unlikely and thus we can consider *ζ*_11_ = *ζ*_22_ = 0. Therefore, the only possible attachment occurs between complementary oligomers and the hybridization vector becomes a scalar *n*_12_ ≡ *n*. In this case, we thus re-obtain the onedimensional theory given by Eq. 6, but with the degeneracy being

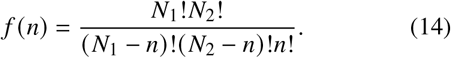

## RESULTS

From now on, we will focus on the two cases introduced above. Namely, the self-complementary (SC) system and the complementary couple (CC), neglecting self-hybridization in the latter. Firstly, let us define the melting curve as the fraction of free oligomers corresponding to the less populated sequence. Therefore, the melting curve is normalized with the maximum number of duplexes. Taking into account that the maximum number of duplexes for the SC or CC cases are *N /*2 (we assume an even *N* for simplicity) or *N_m_* = min *N*_1_, *N*_2_ respectively, we get

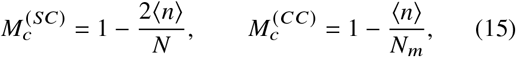

where we have introduced explicitly in the notation if we are referring to SC or to CC. Hence, once the system is defined, that is, the oligomer concentration *c* and the enthalpic Δ*H* and entropic Δ*S* changes of duplex formation are known, the melting curve *M_c_* is just a function of the temperature. On the one hand, for very low temperatures the state with minimum enthalpy, which corresponds with the maximum number of duplexes and then *M_c_* ~ 0, is promoted. On the other hand, entropy is favored for high temperatures and high values of the melting curve *M_c_* ~ 1 are expected.

In the following, we compute the melting curve for our two SC and CC mixtures of oligomers. We provide the results both exactly (for finite systems) and in the thermodynamic limit.

### Exact results for finite systems

Herein, we compute exactly the partition function and the melting curve for SC and CC systems. In order to obtain the partition function, we just need to carry out the sum in Eq. 6 with the corresponding degeneracy (either Eq. 7 for the SC case or Eq. 14 for the CC case). Taking into account that the upper bounds of the sum are, respectively *N*/2 and *N_m_* = min *N*_1_, *N*_2_, and defining conveniently *N_M_* = max *N*_1_, *N*_2_, we get

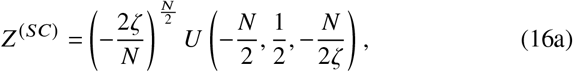

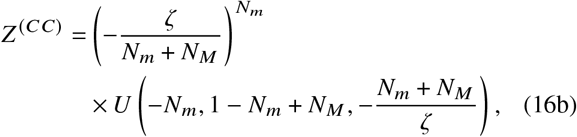

where *U(a*, *b*, *x*) is the confluent hypergeometric function of the second kind also known as Tricomi’s function (32). Performing the derivative in Eq. 8 and introducing it in Eq. 15 yields

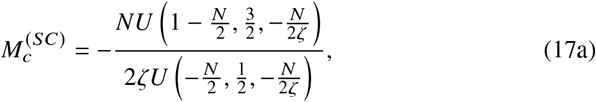

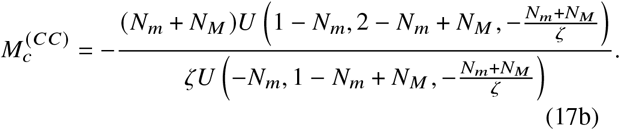

Remarkably, the expressions above are exact. The prediction should be valid even when the mixture has very few component. Of course, real experimental conditions imply *N* ⪢ 1. Moreover, we expect from the physical point of view that in such a limit, at fixed number concentration *c* = *N/V*, *N* does not play any significant role as shown in the first three panels of Fig. 1.

**Figure 1:**
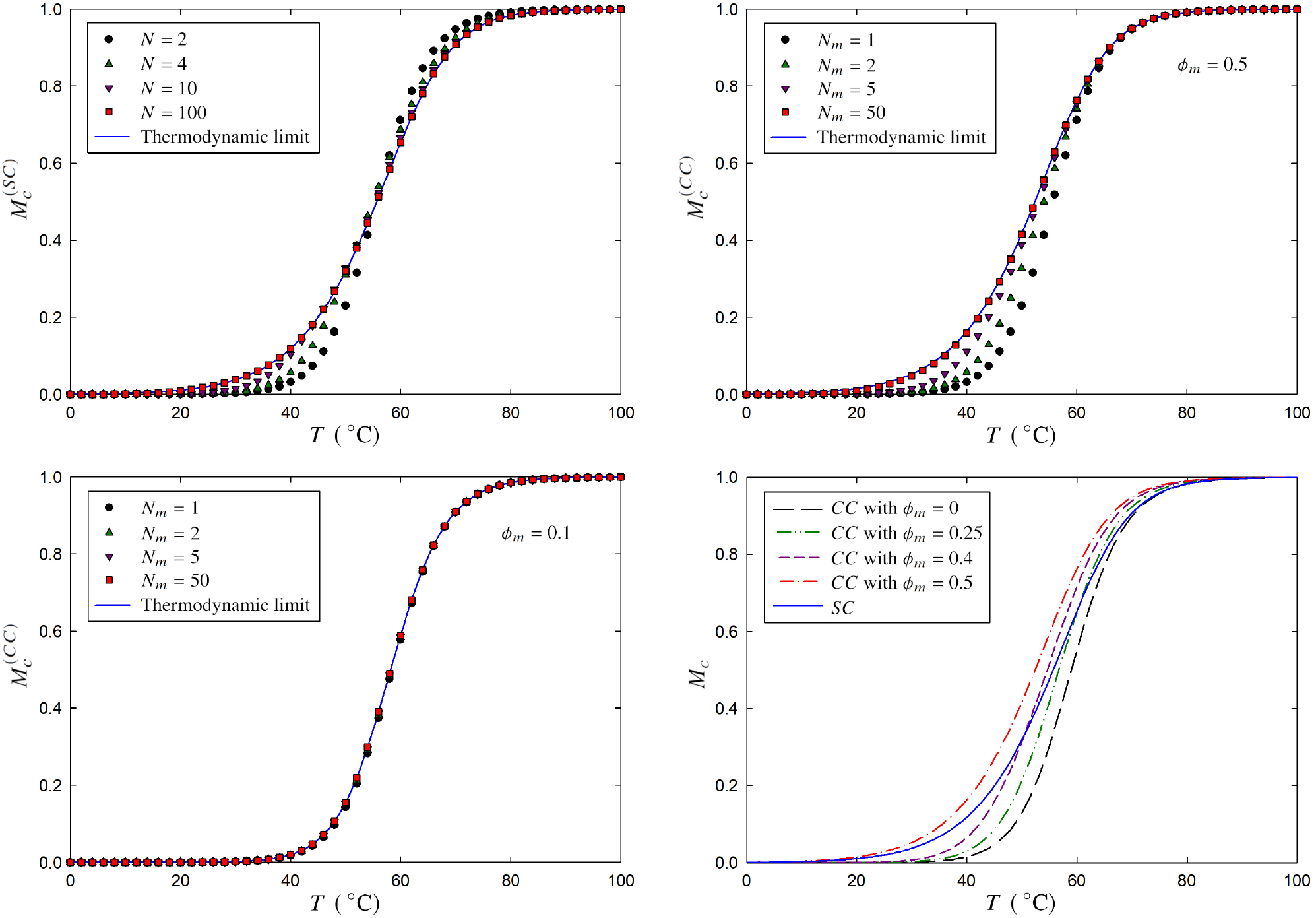
Melting curve for different DNA mixtures. The convergence of the melting curves for finite systems to the predicted thermodynamic limit when the number of constituents is increased is shown in a SC system (top left), a balanced CC system with *ϕ_m_* = 0.5 (top right), and an unbalanced CC system with *ϕ_m_* = 0.1 (bottom left). Bottom right panel presents a comparison of thermodynamic melting curves for CC systems with different values of *ϕ_m_* and a SC system. For the numeric evaluation, we have assumed Δ*H* = −45 kcal · mol^−1^ = −120 cal · mol^−1^ · K^−1^ and a concentration of *c* = 0.4 mmol/L.

### Thermodynamic limit

Since reasonable experimental conditions assure that *N* ⪢ 1, we focus herein on studying in detail this limit. Furthermore, depending on the arguments, hypergeometric functions are not always easy to compute, which makes even more evident the appropriateness of a more illuminating approach. We consider the thermodynamic limit, which entails the limit *N* → ∞ but with the concentration *c* = *N /V* remaining fixed.

Since performing asymptotic study of the ratios of hypergeometric functions in Eq. 17 is not an easy task, we approach the limit in an alternative way. Specifically, we use Grassmann variables (33) that allow us to derive the partition function in the thermodynamic limit through saddle point integration (34) after a Hubbard–Stratonovich transformation (35). Doing so, we are able to compute the melting curve in the thermodynamic limit for both systems of interest, resulting in

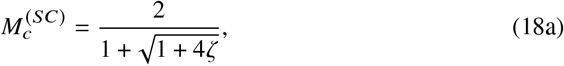

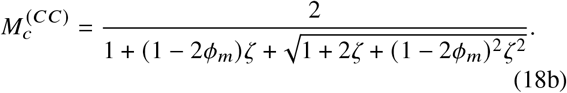

where we have introduced the fraction of the less concentrated sequence *ϕ_m_* = *N_m_*/*N*. The details of the calculations are provided in the Supporting Material. The convergence of the melting curve to the predicted thermodynamic limit is shown in the three first panels of Fig. 1

### Melting temperature

In the context of DNA hybridization and melting, the most analyzed quantity is the melting temperature *T_m_*. The melting temperature is simply defined as the temperature in which half of duplexes are formed. According to our definition of melting curve, the melting temperature is obtained when we equal *M_c_* = 0.5, resulting in

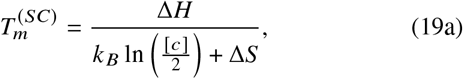

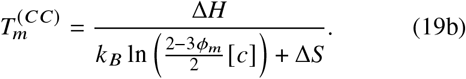

## DISCUSSION

Now, we highlight the most significant facts that stem from the results derived in the previous section. In Fig. 1, we have illustrated several melting curves for both, SC and CC, cases either in systems with finite size or in the thermodynamic limit. Specifically, we can observe how the convergence to the thermodynamic limit, when increasing the number of constituents, is perfect in the first three panels. Remarkably, as seen in the bottom left panel of 1, the effect of the finite size almost disappears for very unbalanced CC systems, that is, with low values of *ϕ_m_*. This is in fact intuitive since, in this limit, the less concentrated sequence can be considered as in contact with a bath of strands of the more concentrated sequence. Therefore, the finite size becomes irrelevant since the population of the more concentrated sequence takes relatively high values, legitimating the thermodynamic result, even for low values of *N_m_*. In other words, if we consider a very unbalanced case, that is, *ϕ* ⪡ 0.5, although *N_m_* could be low, *N* will take high values. Finally, in the bottom right panel, we have plotted several melting curves in the thermodynamic limit of CC systems for different fractions of the less concentrated sequence *ϕ_m_*, as well as the thermodynamic limit of an equivalent SC system. For all the numerical evaluations of the melting curves, we have assumed Δ*H* = −45 kcal · mol^−1^, Δ*S* = −120 cal · mol^−1^ · K^−1^ and a total concentration of *c* = 0.4 mmol/L. The thermodynamic parameters considered are consistent with the order of magnitude of complementary sequences around a length of 6 bases at a reference salt concentration of 1*M* (26).

### Exact results for finite systems

Our exact theory culminating in Eq. 17 provides an analytical prediction for the melting curve in systems with finite size either SC or CC. Considering equivalent free energies and concentrations, the SC system will be more stable than the balanced CC case. In a SC system, all components are available for hybridization, whereas the identity of the oligomer in the CC system is important. Nevertheless, there is a curious unique situation where both of them converge to the same phenomenology. This is when we compare a SC system with *N* = 2 and a *CC* system with *N*_1_ = *N*_2_ = 1. Under this assumption, the partition function in both cases becomes simply

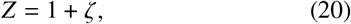

thus obtaining the melting curve

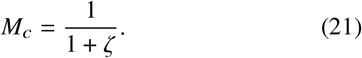

Our theory, as it must be, reflects this consistency. In particular, the melting curves plotted with black circles in the top panels of Fig. 1 are completely identical.

For the best of our knowledge, it is the first time that exact results, beyond the thermodynamic limit, are derived in the context of DNA melting. Nevertheless, as stated in the Introduction, the problem of DNA hybridization resembles the (dis)association problem in the field of chemical reactions. Specifically, the SC and CC case analyzed in this work have a direct parallelism with the formation of diatomic molecules either homonuclear or heteronuclear respectively. In spite of a different approach, based on computing the stationary solution of the Master Equation describing the dynamics of the reaction, the results derived by McQuarry (30) in the context of chemical reactions are fully consistent with ours.

### Thermodynamic limit

The convergence of the exact result for increasing values of *N* to the derived thermodynamic limit shown in the first three panels of Fig. 1 represents a direct validation of our asymptotic calculations. On the one hand, the result for the melting curve in the case of CC systems matches with those provided in the literature. Specifically, most of the time, one finds the results derived in the particular case of a balanced CC system, with *ϕ_m_* = 0.5. Our result goes beyond the balance case of CC mixtures, explaining the role of the relative concentration through the fraction *ϕ_m_*. This is especially clear in the bottom right panel of Fig. 1, where we have displayed the melting curves for the CC system using different values of *ϕ_m_* as well as we compare with the melting curve for the SC system. Therein, we observe, as stated above, that the SC system is more stable than the balanced CC case, that is, the melting curve for the first remains below the one for the latter. Furthermore, we can see how the melting curve for *ϕ_m_* = 0 acts as a lower bound for the SC case. Note that when we write *ϕ_m_* = 0, we are considering the limit of *ϕ_m_* → 0 after taking the thermodynamic limit. Indeed, in this extremely unbalanced case there is a bath of available strands to be paired. On the other hand our result for the melting curve for the SC case differs from those traditionally found in the literature (28). Therein, one finds a factor 8 accompanying *ζ* in the denominator instead of the factor 4 written in Eq. 18.

Such a mismatch is quite curious. On the one hand, our approach (with the 4) is the correct asymptotic limit of the exact results that are consistent with the theory developed for chemical reactions. On the other hand, the traditional melting curve in the literature (with the 8) has been used intensively explaining and fitting data from real experiments. This leads to the question: what is special in self-complementary sequences?

From the physical point of view, a duplex is characterized by its Δ*G*, which, apart from the temperature, is defined by its enthalpy Δ*H* and its entropy Δ*S*. For both cases, either SC or CC, there is just a single way for the coupling of two oligomers, due to the directionality 5’ – 3’ of the single strand DNA. Therefore, there is no physical intuition for stating that the biophysical mechanisms of hybridization are different when the oligomers are self-complementary. However, in the tables of thermodynamic parameters for SC and CC present in literature, an entropic *symmetry correction* is systematically applied to self-complementary sequences (19, 21, 22, 25, 26, 28). This correction Δ*S_sym_* is such that exp (Δ*S_sym_/k_B_*) ≃ 0.5 In other words, this artificial correction transforms the *wrong* factor 8 into the *correct* one 4.

Of course, from the practical point of view, there is no difference between using the traditional melting curve after applying the symmetric correction or using directly our result without adding any artifact. Nevertheless, from our point of view, it is important to highlight that there is neither a real extra entropic contribution nor a special physics in self-complementary DNA.

### Reaction equations approach

The traditional result for the SC case has been usually derived from equilibrium constant arguments. Herein, we give a brief alternative approach which highlight where the mismatch emerges. Specifically, we consider the reaction

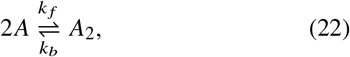

which represents the (dis)association of homonuclear diatomic molecules, which also describes the hybridization of self-complementary systems. We write down the macroscopic evolution equation, that is, neglecting fluctuations, for the number of free atoms *N_a_*,

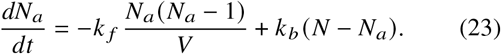

In the equation above, accompanying the forward rate *k_f_*, we find the number of possible couples in the system normalized by the volume which is *N_a_ (N_a_* – 1)/2*V* that multiplies the number of atoms that disappear, that is, 2. In turn, along with the backward rate *k_b_*, we have the number of duplexes *N_a_ (N_a_* – 1)/2 that multiplies the number of appearing atoms, which is again 2.

The equilibrium constant, defined as the ratio of the forward rate, yields *K* = *k_f_ k_b_* = exp(*β*Δ*G*). Introducing this relation, the definition of the melting curve *M_c_* = *N_a_ N* and the stationary solution of Eq. 23, then the relation 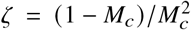 is obtained, where we have made use of the definition of *ζ* in Eq. 4 and neglected terms of order *N*^−1^. Solving this equation for *M_c_* leads straightforwardly to our result for the SC melting curve in Eq. 18. In the traditional derivation, the mismatch stems from taking blindly the equilibrium constant as the fraction between the concentration of duplexes and the squared concentration of the single strands. Such misconception is equivalent to assume in our equations that the forward process is proportional to the product 2*N_a_* (*N_a_ –* 1)/*V* instead of *N_a_* (*N_a_ –* 1)/*V*. This is not correct since it means that the possible couples of free particle are counted twice.

## CONCLUSION

In this work, we have put forward the fundamentals of statistical physics to the service of the problem of DNA melting. We have provided an equilibrium description of hybridization and melting for nucleic acids, deriving the partition function in any arbitrary mixture made by DNA.

We have then focused on two experimentally relevant systems, made by either self-complementary sequences (SC) or different complementary couple (CC) of strands. In such cases, the exact melting curves have been derived and we have shown that our results agree with the known expressions in the context of chemical reactions. Remarkably, this implies that there is a factor two mismatch with the literature of DNA melting. Such a discrepancy has led us to debunk the artificial entropic correction applied to self-complementary duplexes in the literature, which we have now clearly identified as an artifact performed in order to reconcile experimental data with the theory.

Beyond our exact result, we have derived the melting curve of the systems of interest in the thermodynamic limit, which is the meaningful situation in real experimental conditions. As expected, the convergence to the thermodynamic limit when considering the exact result with increasing number of constituents is excellent.

Although the computation of the melting curve for the SC and CC cases has been made by considering only complementary sequences (and thus just one single way for the attachment between pairs of DNA strands), our model can be generalized to describe pairs with a preferential ways of attachment, even if not complementary. The lack of full complementarity would lead to lower values of *ζ*, resulting in a system with lower stability and a melting curve shifted to lower temperatures. The exact calculations for such more heterogeneous cases is left for future investigations.

## AUTHOR CONTRIBUTIONS

S.S. and T.B. designed the research. A.M. provided the original idea for the analytical approach. C.A.P. and S.M. performed the research. All authors contributed to the manuscript writing.

## ACKNOWLEDGMENTS

S.S. and C.A.P acknowledge UNIPD STARS grant BioReact 2018. T.B. acknowledges support by PRIN2017 project from Ministero Istruzione, Università e Ricerca [ID 2017Z55KCW]. A. Maritan acknowledges the support from University of Padova through “Excellence Project 2018” of the Cariparo foundation

## SUPPORTING MATERIAL

An online supplement to this article can be found by visiting BJ Online at http://www.biophysj.org.

## REFERENCES

1. Watson, J., and F. Crick, 1953. Molecular Structure of Nucleic Acids: A Structure for Deoxyribose Nucleic Acid. Nature 171:737–738. https://doi.org/10.1038/171737a0.

2. Ririe, K. M., R. P. Rasmussen, and C. T. Wittwer, 1997. Product Differentiation by Analysis of DNA Melting Curves during the Polymerase Chain Reaction. Analytical Biochemistry 245:154–160. http://www.sciencedirect.com/science/article/pii/S0003269796999169.

3. Reed, G. H., J. O. Kent, and C. T. Wittwer, 2007. High-resolution DNA melting analysis for simple and efficient molecular diagnostics. Pharmacogenomics 8:597–608. https://doi.org/10.2217/14622416.8.6.597, pMID: 17559349.

4. Cuesta-Lopez, S., D. Angelov, and M. Peyrard, 2009. Adding a new dimension to DNA melting curves. EPL (Europhysics Letters) 87:48009. https://doi.org/10.1209%2F0295-5075%2F87%2F48009.

5. Von Keyserling, H., T. Bergmann, M. Wiesel, and A. M. Kaufmann, 2011. The use of melting curves as a novel approach for validation of real-time PCR instruments. BioTechniques 51:179–184. https://www.future-science.com/doi/abs/10.2144/000113735, pMID: 21906039.

6. Balog, J. A., L. Z. Fehér, and L. G. Puskás, 2017. Decoding DNA labels by melting curve analysis using real-time PCR. BioTechniques 63:261–266. https://www.future-science.com/doi/abs/10.2144/000114618, pMID: 29235972.

7. Mandel, M., and J. Marmur, 1968. [109] Use of ultraviolet absorbance-temperature profile for determining the guanine plus cytosine content of DNA. In Nucleic Acids, Part B, Academic Press, volume 12 of Methods in Enzymology, 195–206. http://www.sciencedirect.com/science/article/pii/0076687967121332.

8. Wells, R., J. Larson, R. Grant, B. Shortle, and C. Cantor, 1970. Physicochemical studies on polydeoxyribonucleotides containing defined repeating nucleotide sequences. Journal of Molecular Biology 54:465–497. http://www.sciencedirect.com/science/article/pii/002228367090121X.

9. Wartell, R. M., and A. S. Benight, 1985. Thermal denaturation of DNA molecules: A comparison of theory with experiment. Physics Reports 126:67–107. http://www.sciencedirect.com/science/article/pii/0370157385900602.

10. Mergny, J.-L., and L. Lacroix, 2003. Analysis of Thermal Melting Curves. Oligonucleotides 13:515–537. https://doi.org/10.1089/154545703322860825, pMID: 15025917.

11. Owczarzy, R., 2005. Melting temperatures of nucleic acids: Discrepancies in analysis. Biophysical Chemistry 117:207–215. http://www.sciencedirect.com/science/article/pii/S0301462205001201.

12. Azbel, M. Y., 1979. Phase transitions in DNA. Phys. Rev. A 20:1671–1684. https://link.aps.org/doi/10.1103/PhysRevA.20.1671.

13. Dauxois, T., M. Peyrard, and A. R. Bishop, 1993. Dynamics and thermodynamics of a nonlinear model for DNA denaturation. Phys. Rev. E 47:684–695. https://link.aps.org/doi/10.1103/PhysRevE.47.684.

14. Tinoco, I., O. C. Uhlenbeck, and M. D. Levine, 1971. Estimation of Secondary Structure in Ribonucleic Acids. Nature 230:362–367. https://doi.org/10.1038/230362a0.

15. Borer, P. N., B. Dengler, I. Tinoco, and O. C. Uhlenbeck, 1974. Stability of ribonucleic acid doublestranded helices. Journal of Molecular Biology 86:843–853. http://www.sciencedirect.com/science/article/pii/002228367490357X.

16. Ghosh, S., S. Takahashi, T. Endoh, H. Tateishi-Karimata, S. Hazra, and N. Sugimoto, 2019. Validation of the nearest-neighbor model for Watson-Crick self-complementary DNA duplexes in molecular crowding condition. Nucleic Acids Research 47:3284–3294. https://doi.org/10.1093/nar/gkz071.

17. Marky, L. A., and K. J. Breslauer, 1982. Calorimetric determination of base-stacking enthalpies in double-helical DNA molecules. Biopolymers 21:2185–2194. https://onlinelibrary.wiley.com/doi/abs/10.1002/bip.360211107.

18. Petersheim, M., and D. H. Turner, 1983. Base-stacking and base-pairing contributions to helix stability: thermodynamics of double-helix formation with CCGG, CCGGp, CCGGAp, ACCGGp, CCGGUp, and ACCGGUp. Biochemistry 22:256–263. https://doi.org/10.1021/bi00271a004, pMID: 6824629.

19. Breslauer, K. J., R. Frank, H. Blöcker, and L. A. Marky, 1986. Predicting DNA duplex stability from the base sequence. Proceedings of the National Academy of Sciences 83:3746–3750. https://www.pnas.org/content/83/11/3746.

20. Marky, L. A., and K. J. Breslauer, 1987. Calculating thermodynamic data for transitions of any molecularity from equilibrium melting curves. Biopolymers 26:1601–1620. https://onlinelibrary.wiley.com/doi/abs/10.1002/bip.360260911.

21. Serra, M. J., and D. H. Turner, 1995. [11] Predicting thermodynamic properties of RNA. In Energetics of Biological Macromolecules, Academic Press, volume 259 of Methods in Enzymology, 242–261. http://www.sciencedirect.com/science/article/pii/0076687995590471.

22. Sugimoto, N., S.-i. Nakano, M. Yoneyama, and K.-i. Honda, 1996. Improved Thermodynamic Parameters and Helix Initiation Factor to Predict Stability of DNA Duplexes. Nucleic Acids Research 24:4501–4505. https://doi.org/10.1093/nar/24.22.4501.

23. Gray, D. M., 1997. Derivation of nearest-neighbor properties from data on nucleic acid oligomers. I. Simple sets of independent sequences and the influence of absent nearest neighbors. Biopolymers 42:783–793.

24. Allawi, H. T., and J. SantaLucia, 1997. Thermodynamics and NMR of Internal G·T Mismatches in DNA. Biochemistry 36:10581–10594. https://doi.org/10.1021/bi962590c, pMID: 9265640.

25. SantaLucia, J., 1998. A unified view of polymer, dumbbell, and oligonucleotide DNA nearest-neighbor thermodynamics. Proceedings of the National Academy of Sciences 95:1460–1465. https://www.pnas.org/content/95/4/1460.

26. SantaLucia, J., and D. Hicks, 2004. The Thermodynamics of DNA Structural Motifs. Annual Review of Biophysics and Biomolecular Structure 33:415–440. https://doi.org/10.1146/annurev.biophys.32.110601.141800, pMID: 15139820.

27. Benight, A. S., and R. M. Wartell, 1983. Influence of base-pair changes and cooperativity parameters on the melting curves of short DNAs. Biopolymers 22:1409–1425. https://onlinelibrary.wiley.com/doi/abs/10.1002/bip.360220512.

28. Owczarzy, R., P. M. Vallone, F. J. Gallo, T. M. Paner, M. J. Lane, and A. S. Benight, 1997. Predicting sequence-dependent melting stability of short duplex DNA oligomers. Biopolymers 44:217–239.

29. Dimitrov, R. A., and M. Zuker, 2004. Prediction of Hybridization and Melting for DoubleStranded Nucleic Acids. Biophysical Journal 87:215–226. http://www.sciencedirect.com/science/article/pii/S0006349504735110.

30. McQuarrie, D. A., 1967. Stochastic approach to chemical kinetics. Journal of Applied Probability 4:413–478.

31. McCoy, B. J., and R. G. Carbonell, 1977. Master equation theory for steady-state chemical reactions: Dissociation of diatomic molecules in gases. The Journal of Chemical Physics 66:4564–4571. https://doi.org/10.1063/1.433712.

32. Lebedev, N., and R. Silverman, 1972. Special Functions and Their Applications. Dover Books on Mathematics. Dover Publications. https://books.google.it/books?id=po-6Yxz851MC.

33. Berezin, F. A., N. Mugibayashi, and A. Jeffrey, 1966. The Method of Second Quantization. Pure and applied physics: a series of monographs and textbooks. 24. Academic Press. https://books.google.it/books?id=fAlRAAAAMAAJ.

34. Bender, C. M., and S. A. Orszag, 1999. Advanced Mathematical Methods for Scientists and Engineers. Springer, New York.

35. Hubbard, J., 1959. Calculation of Partition Functions. Phys. Rev. Lett. 3:77–78. https://link.aps.org/doi/10.1103/PhysRevLett.3.77.

